# Assembly processes of bacterial and fungal community differ between desert and oasis habitats in an arid inland river basin, northwest China

**DOI:** 10.1101/2023.06.21.545439

**Authors:** Wen-Juan Wang, Yi-Ming Ding, Ming-Xun Ren, Jing-Wen Li

**Author notes:** Corresponding author: School of Ecology and Nature Conservation, Beijing Forestry University, 35 Tsinghua East Road, Haidian District, Beijing 100083, China.

## Abstract

- Oasis habitat play a critical role in arid areas, yet little is known about microbial community assembly processes and its differences in oasis and desert habitat in arid inland river basin.
- Herein, using 21 and 18 sample data respectively collected from oasis and desert habitats at the lower reaches of Heihe River, northwest China, we examined the assembly processes of soil bacterial and fungal communities and revealed the driving factors affecting the relative contributions of deterministic and stochastic processes.
- We found that deterministic processes, especially variable selection, dominated bacterial community assembly in oasis habitat, while stochastic processes were more important in desert habitat. By contrast, stochastic processes dominated fungal community assembly both in oasis and desert habitats, in which dispersal limitation played a more important role. Environmental (e.g. EC) and biotic factors (microbial species associations) significantly mediated the bacterial community assembly. However, both environmental and biotic factors had little/no effects on fungal community assembly.
- This study highlights the soil microbial community assembly is habitat- and taxon-dependent, and environmental (EC) and biotic factors play an important role in regulating these assembly processes in arid inland river basin.

## Introduction

Soil microorganisms play important roles in ecosystem functions, such as biogeochemical cycle (Bardgett and van der Putten, 2014; Sokol *et al*., 2022), verifying their distribution patterns and assembly mechanisms can promote our understanding of the relationship between soil microbial communities and ecosystem functions (Hanson *et al*., 2012; Thakur *et al*., 2020; Jiao *et al*., 2022). Bacteria and fungi are two major taxa of microorganisms, which harbored different morphological, physiological and ecological characteristics (Hannula *et al*., 2017). For instance, fungal “body size” is usually larger than bacteria (Chen *et al*., 2020). In addition, fungal species are often identified as the primary decomposers of recalcitrant organic materials (van den Brink and de Vries, 2011; van der Wal *et al*., 2013), such as cellulose and lignin, while bacteria utilize labile organic matter and simultaneously decompose dead fungal mycelia (Rousk *et al*., 2016; Llado *et al*., 2017). These differences would arise different distribution patterns and assembly processes of bacterial and fungal communities (Wang *et al*., 2020). Indeed, some previous studies reported that bacterial and fungal communities distributed differently and were driven by different factors (Wang *et al*., 2017; Bahram *et al*., 2021). Studies in a typical dryland ecosystem showed that soil moisture and pH explained more variations of bacterial structure, while geographic distance and aridity index were more important factors for fungal community variations (Wang *et al*., 2017). However, it is still few to simultaneously study assembly processes of bacterial and fungal communities.

Assembly processes fall into two categories, namely deterministic processes based on the niche theory and stochastic processes based on the neutral theory (Dumbrell *et al*., 2010). The former emphasizes the role of environmental filtering and interspecific interactions in existence, relative abundance and turnover of microorganisms (Vellend, 2010), while the latter emphasizes the role of dispersal and ecological drift in shaping distribution patterns of microbial communities (Zhou and Ning, 2017). Hitherto, we have reached a general consensus that deterministic and stochastic processes simultaneously determined assembly of microbial communities (Stegen *et al*., 2012; Wang *et al*., 2013). However, the relative contributions of these two processes could be varied with our targeting ecosystems and/or microbial types, which was mediated by a variety of factors (Jiao and Lu, 2020; Ni *et al*., 2021; Yang *et al*., 2022). For example, stochasticity rather than determinism exerted a greater impact on the archaeal community assembly in coastal sediments (Liu *et al*., 2020), whereas determinism was more important in microbial (bacteria and archaea) community assembly that strong selection pressure was imposed by salinity in desert ecosystem (Zhang *et al*., 2019). Therefore, it is essential to simultaneously explore community assembly processes for different microbial types and different habitats.

Within the framework of deterministic and stochastic processes, influence of environmental factors on microbial community assembly processes have been studied along a wide range of environments or habitats (Tripathi *et al*., 2018; Barnett *et al*., 2020). However, far fewer studies explore the roles of biotic interactions in shaping community assembly, which could determine the microbial community structure by competition and mutualisms (Zhang *et al*., 2018). Species co-occurrence networks and their network topological features are appropriate methods to infer the biotic interactions (Deng *et al*., 2012; Ma *et al*., 2020). Based on the topological features, we could fill the gap in understanding how biotic interactions affected on microbial community assembly and how environmental factors affected biotic interactions and indirectly influenced microbial community assembly.

The arid inland river basin is popularly known as a “hot spot” for biodiversity due to its higher water availability in arid and semi-arid regions (Hao *et al*., 2010). Meanwhile, different water conditions also facilitate the formation of the two obviously different habitats, namely oasis and desert, in arid regions (Li *et al*., 2016; Wang *et al*., 2023). Our previous studies have proved that composition and distribution patterns of microbial communities and its driving factors were significantly different in these two habitats (Wang *et al*., 2019; Wang *et al*., 2021a). However, it is unclear whether assembly processes of bacterial and fungal communities are different in these two habitats. Generally, stronger deterministic processes were observed on larger scales due to extensive environment gradients, while stochastic processes occupied a larger proportion on small scales (Zhao *et al*., 2019a; Wang *et al*., 2020). Ejina oasis, located in arid regions of northwest China and shaped by the lower reaches of Heihe River, harbored greater environmental heterogeneity in the small geographic area (Wang, *et al*., 2019; Zhao *et al*., 2019b). However, relatively little is known how deterministic and stochastic processes is balanced in such small but high environment heterogeneity areas.

In this study, we aim to addressed the following questions: 1) whether soil bacterial and fungal community assembly processes are different in oasis and desert habitats? 2) How do environmental factors and biotic interactions mediate the relative importance of deterministic and stochastic processes in bacterial and fungal community assembly? These questions could be well addressed based on the existing high-throughput sequencing data sets obtained from the oasis and desert habitats at the lower reaches of Heihe River, northwest China (Wang *et al*., 2019; Wang *et al*., 2021a).

## Materials and methods

### Data sets

We used high-throughput sequencing, soil properties and plant attributes data from studies of soil bacterial and fungal diversity in oasis and desert habitats (Wang *et al*., 2019; Wang *et al*., 2021a). The sample site was located in the lower reaches of the Heihe River, northwest China, and the methods of field sampling had been described in previous study (Wang *et al*., 2019). Briefly, 21 and 18 samples were respectively collected from the oasis and desert habitats. Soil properties, including available nitrogen (AN), soil electrical conductivity (EC), soil particle composition (sand, silt, and clay content percentage), etc. and plant attributes, including plant community, specific leaf area (SLA), leaf nitrogen content (LNC), etc. were determined using standard methods (Wang *et al*., 2019). Sequence data of the V3-V4 regions for bacterial communities and the ITS regions for fungal communities were acquired from an Illumina Miseq platform (Allwegene Technology, Beijing, China). Sequences were filtered for quality control and classified into OTUs based on a 97% threshold. And each OTU was assigned to taxonomic groups through the RDP Classifier against the Silva128 16S rRNA database for bacteria, and the UNITE database for fungi.

### Co-occurrence network analysis

To estimate effects of biotic interactions on microbial community assembly, cross-kingdom co-occurrence networks consisting of bacterial and fungal taxa was constructed. To reduce rare OTUs in the data sets, we selected OTUs for Spearman correlation calculation based on the following criteria. Firstly, we selected phyla with a mean relative abundance greater than 1%, and then OTUs with relative abundance greater than 0.01%, which were simultaneously present in more than 20% of all soil samples, were retained (Liu *et al*., 2020). Robust correlations with Spearman’s correlation coefficients |ρ| > 0.7 and p < 0.001 were used to construct networks with “igraph” package, while all p-values were adjusted using Benjamini-Hochberg standard false discovery rate correction (FDR-BH) (Benjamini *et al*., 2006). It was visualized using Gephi (http://gephi.github.io/).

Then, we extracted sub-network by preserving the OTUs of individual soil samples using the induced_subgraph function in “igraph” package in R (Csardi and Nepusz, 2006). The topology features of each sub-network, including average clustering coefficient (ACC), average degree (AD), average path length (APL), the proportion of interacted associations between bacterial and fungal taxa (Bac_Fun), density of the network (Den), modularity (Mod), and the proportion of negative associations (Neg), were calculated to estimate the potential biotic associations (Jiao *et al*., 2022).

### Community assembly analysis

An important assumption is significant phylogenetic signal when we evaluate the microbial community assembly processes based on phylogenetic turnover between communities (Dini-Andreote *et al*., 2015). To test this assumption, we firstly calculated the relative-abundance-weighted mean value of each soil variable for each OTU as its environmental optima (Wang *et al*., 2013; Dini-Andreote *et al*., 2015). Then, we calculated the correlation coefficients between niche differences (pairwise Euclidean distances of OTUs’ environmental optima) and phylogenetic distance under different phylogenetic distance thresholds using Mantel correlogram function in “vegan” package (Oksanen *et al*., 2022). These correlation coefficients were tested by p-values calculated based on 999 permutations of the distance matrix with Bonferroni correction (Dini-Andreote *et al*., 2015). The result was consistent with previous reports that showed significant phylogenetic signals at relatively short phylogenetic distances (Fig. S1), supporting the assumption of this study(Wang *et al*., 2013; Liu *et al*., 2020).

Pairwise phylogenetic turnover between communities (βMNTD) were calculated using comdistnt function in “picante” package in R (Kembel *et al*., 2010). Then, a null model analysis was used to evaluate assembly processes of bacterial and fungal communities (Stegen *et al*., 2013). βNTI (variations in phylogenetic turnover (βMNTD)) were measured by null model, which quantified the magnitude and direction of deviations of observed βMNTD from mean of the null model-based βMNTD distribution. βNTI values > +2 means significantly greater phylogenetic turnover than expected, indicating variable selection; meanwhile, βNTI values < −2 means significantly smaller phylogenetic turnover than expected, indicating homogeneous selection. |βNTI| < 2 means that phylogenetic turnover is not significantly different from expectation of null model, emphasizing the importance of stochastic processes on community turnover. Further, RC_bray_ metric, the standardized taxonomic β-diversity, was used to differentiate stochastic processes into dispersal limitation, ecological drift and homogenizing dispersal (Stegen *et al*., 2013). Specifically, the relative contribution of dispersal limitation was estimated by calculating the fraction of pairwise comparisons with |βNTI| < 2 and RC_bray_ > 0.95; the relative contribution of homogenizing dispersal was estimated as the fraction of pairwise comparisons with |βNTI| < 2 and RC_bray_ < −0.95; all fraction of pairwise comparisons with |βNTI| < 2 and |RC_bray_| < 0.95 quantified the contribution of ecological drift (Stegen *et al*., 2013). All estimations above were processed in oasis habitat, desert habitat and the whole samples, respectively.

### Statistical analysis

To access the relative importance of deterministic and stochastic processes in oasis and desert habitats, we compared all βNTI values in these two habitats based on one-way ANOVA. Mantel and partial mantel test were respectively used to examine the influence of soil environmental distance, plant community dissimilarity, spatial distance and plant functional trait distance on bacterial and fungal phylogenetic turnover (βNTI) (Oksanen *et al*., 2022). Then, we further used multiple regression on matrices (MRM) to clarify the explanations of soil properties, plant community and spatial factors on bacterial and fungal βNTI within “ecodist” package (Goslee and Urban, 2007; Lichstein, 2007). To avoid strong collinearity between variables, the varclus function in “Hmisc” package was used to evaluate the redundancy of variables before performing MRM, and the variable of sand content was removed because of Pearson’s ρ^2^ > 0.8 while all the other variables were entered into MRM analysis (Harrell Jr, 2023). In order to avoid effect of data overfitting, we ran the second MRM analysis that was reported in the paper after removing non-significant variables in first MRM (Wang *et al*., 2017). Relative contribution of each variable on bacterial and fungal βNTI was calculated using calc.relimp function in the “relaimpo” package (Grömping, 2006).

The variations of community assembly processes along the major environmental variables (soil EC and silt content) were assessed using regression analysis through comparing the Euclidean distance of the two variables with βNTI, which significance was estimated by performing mantel function in “ecodist” package (Goslee and Urban, 2007). Further, we divided all samples into low EC and high EC groups (Fig. S2) based on an 8 ms/cm threshold (Lin and Bañuelos, 2015), and accessing the relative importance of deterministic and stochastic processes of these two subgroups.

Mantel tests were used to examine relationships between biotic associations and bacterial and fungal phylogenetic turnover (βNTI) (Goslee and Urban, 2007). Additionally, piecewise SEM was performed using the “piecewiseSEM” package (Lefcheck, 2016) to further evaluate the effects of biotic associations and environmental filtering (soil properties, plant community) and their indirect effects on bacterial and fungal phylogenetic turnover (βNTI) in different habitats (Li *et al*., 2022). In the piecewise SEM analysis, we selected topological features of AD, Den, Neg and Bac_fun as biotic association variables based on varclus function in “Hmisc” package (Fig. S3).

## Results

### Relative influence of assembly processes on bacterial and fungal community in oasis and desert habitats

We found deterministic processes play an important role in bacterial community assembly in oasis habitat with |βNTI| > 2 accounted for 63.3%, and which was significantly larger than that in desert habitat (27.5%), indicating the major role of environmental selection in oasis habitat (Fig. 1a). In contrast, stochastic processes dominated fungal community assembly with |βNTI| > 2 accounted for 13.8% and 19.0% respectively in oasis and desert habitat (Fig. 1b). And βNTI value was significantly smaller in oasis than that in desert habitat, indicating environmental selection was more important for fungal community assembly in desert than in oasis habitat (Fig. 1b).

**Figure 1.**
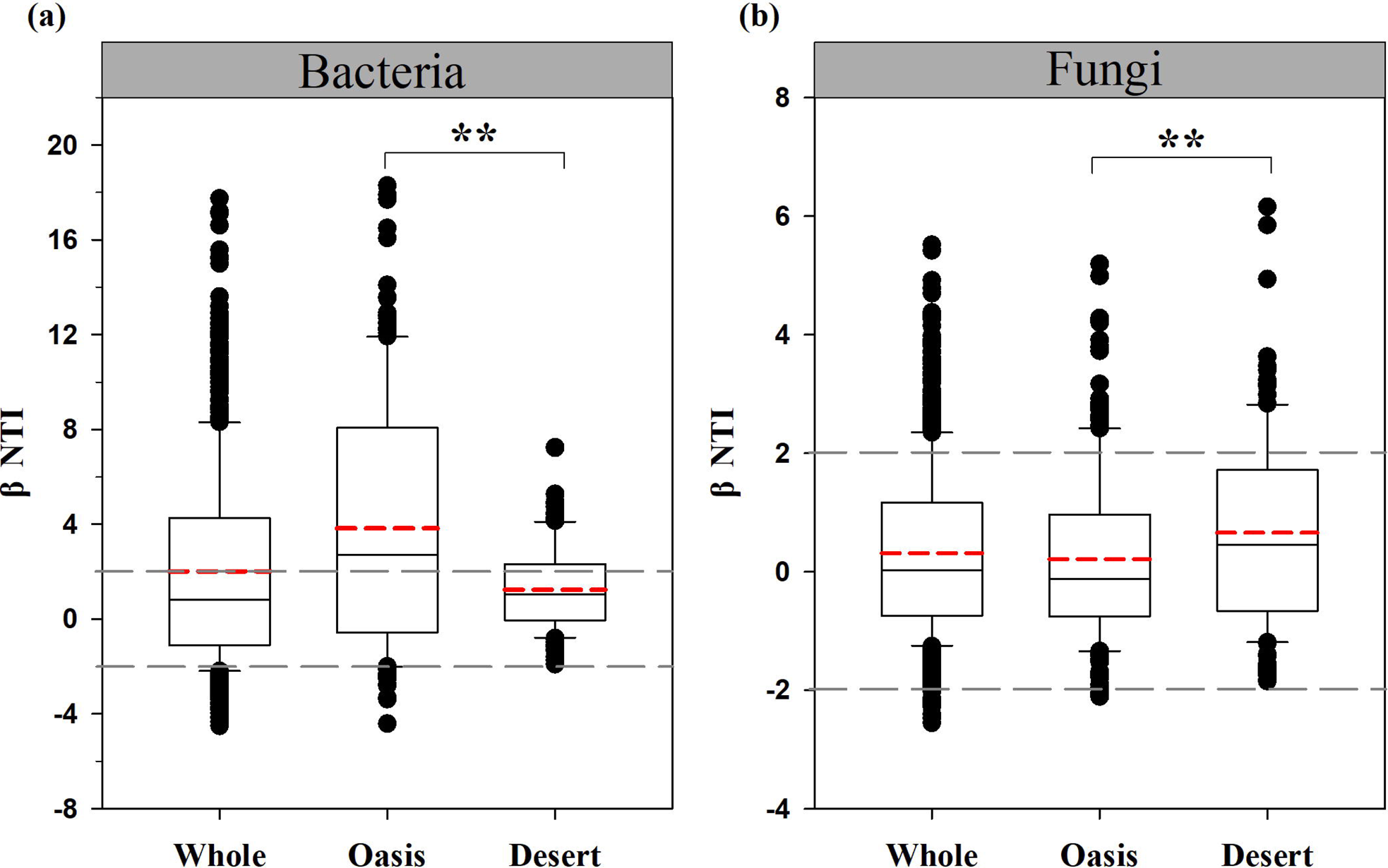
Patterns of βNTI values in different habitats for bacteria (a) and fungi (b). Horizontal dashed gray lines indicate upper and lower significant thresholds at βNTI=+2 and βNTI=-2, respectively. Dashed red lines indicate mean values of βNTI values in different habitats. “**” indicate significant difference of βNTI values at 0.01 level.

Variable selection contributed the largest fraction to bacterial community assembly in the whole samples (39.0%) and oasis habitat (53.3%), and dispersal limitation followed (34.8%, 24.8% respectively), whereas dispersal limitation dominated bacterial community assembly in desert habitat (57.5%) and variable selection followed (27.5%) (Fig. 2a). In contrast, dispersal limitation dominated fungal community assembly, which respectively explained 73.7%, 83.8% and 58.8% in the whole samples, oasis and desert habitat (Fig. 2b).

**Figure 2.**
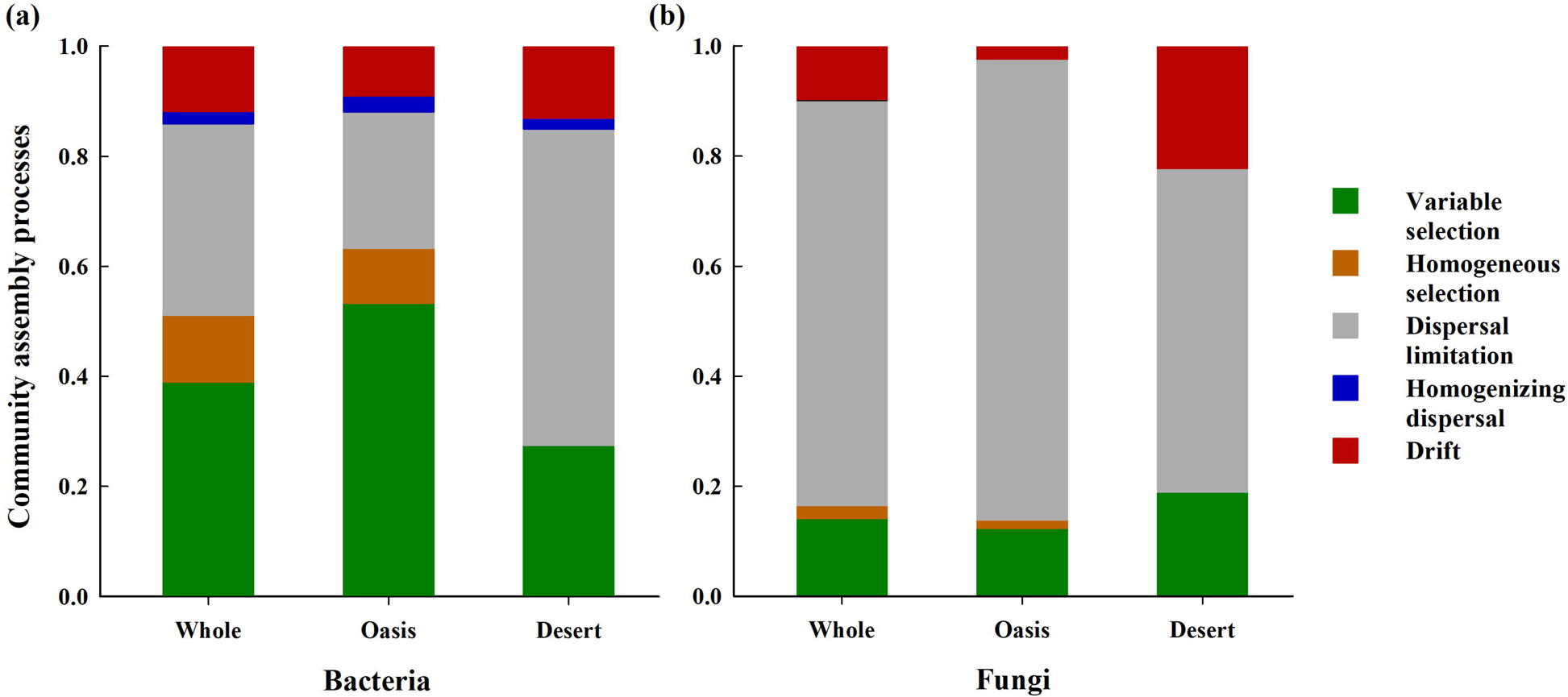
Relative contributions of each ecological assembly process for bacterial (a) and fungal (b) community in different habitats.

### Main factors mediating the balance between deterministic and stochastic processes

Mantel test showed that soil environment and plant community best predicted bacterial community assembly processes, and the relationships remained significant even after controlling for other variables (Table 1). By contrast, fungal community assembly processes had no significant correlations with the investigated variables in oasis habitat whereas it significantly correlated with soil environment in desert habitat (Table 1).

**Table 1.** Mantel and partial mantel test results showing relationships between bacterial and fungal hylogenetic turnover (βNTI) and soil environmental distance, plant community dissimilarity, spatial distance and plant functional trait distance.

|  | Explanatory variables | Controlling for | Whole | Oasis | Desert |
| --- | --- | --- | --- | --- | --- |
| Bacteria | Soil environmental distance (SE) |  | **0.3719\*\*\*** | **0.5473\*\*\*** | **0.2873 \*\*** |
|  | Plant community dissimilarity (PC) |  | **0.1669\*\*** | **0.3766\*\*\*** | **0.2941\*\*** |
|  | Spatial distance (SD) |  | **0.1883\*\*** | 0.0464 | -0.1404 |
|  | Plant functional trait distance (PF) |  | 0.0121 | 0.0974 | 0.0275 |
|  | Soil environmental distance | PC+SD | **0.3302\*\*** | **0.5273\*\*\*** | **0.2437\*** |
|  | Plant community dissimilarity | SE+SD | 0.0733 | **0.3354\*\*** | **0.2777\*\*** |
|  | Spatial distance | SE+PC | 0.0610 | — | — |
| Fungi | Soil environmental distance |  | 0.0663 | 0.0425 | **0.2642\*** |
|  | Plant community dissimilarity |  | 0.0452 | -0.1181 | 0.0008 |
|  | Spatial distance |  | **0.0963\*** | 0.0267 | 0.1020 |
|  | Plant functional trait distance |  | -0.1022 | -0.1386 | -0.1051 |

Note: \*\*\* $p < 0.001$ ; \*\* $p < 0.01$ ; \* $p < 0.05$ .

Results of MRM analysis showed that EC (14.1%) and soil silt content (16.3%) explained more variation of bacterial βNTI in the whole samples (Table 2). Separately, bacterial βNTI was mainly driven by EC (25.4%) and soil particle composition (37.7%) in oasis habitat while that was mainly driven by EC (17.8%) in desert habitat (Table 2). However, no predictive variable was maintained in MRM analysis for fungal βNTI in the whole samples and in oasis habitat (Table 2), which was consistent with more important role of stochastic processes (>80%) in fungal community assembly. Though EC was retained in the final MRM analysis of fungal βNTI in desert habitat, it only explained 10% variation of fungal βNTI (Table 2).

**Table 2.** The results of multiple regression on distance matrices (MRM) showing relative importance of soil variables, plant community and spatial distance for explaining bacterial and fungal βNTI in Whole, Oasis and Desert region.

|  |  | Variables retained in the model and its individual contribution (%) | R<sup>2</sup> | P value |
| --- | --- | --- | --- | --- |
| Bacteria | Whole | spatial distance (3.1), EC (14.1), silt (16.3) | 0.335 | <0.0001 |
|  | Oasis | plant community (6.5), AN (2.3), EC (25.4), silt (13.7), clay (24.0) | 0.719 | <0.0001 |
|  | Desert | plant community (5.7), EC (17.8) | 0.235 | <0.0001 |
| Fungi | Whole | no variable retained in the final model |  |  |
|  | Oasis | no variable retained in the final model |  |  |
|  | Desert | EC (10) | 0.100 | <0.01 |

The bacterial βNTI in oasis habitat showed significantly and positively correlations with ΔEC and Δsilt, respectively (Fig. 3d,e), which indicating that increasing divergence of soil EC and silt content would lead to a shift of assembly processes of bacterial community from stochasticity to variable selection. However, though fungal βNTI was significantly correlated with ΔEC in desert habitat, the divergence of EC was not enough to bring changes of fungal community assembly processes (Fig. 3i).

**Figure 3.**
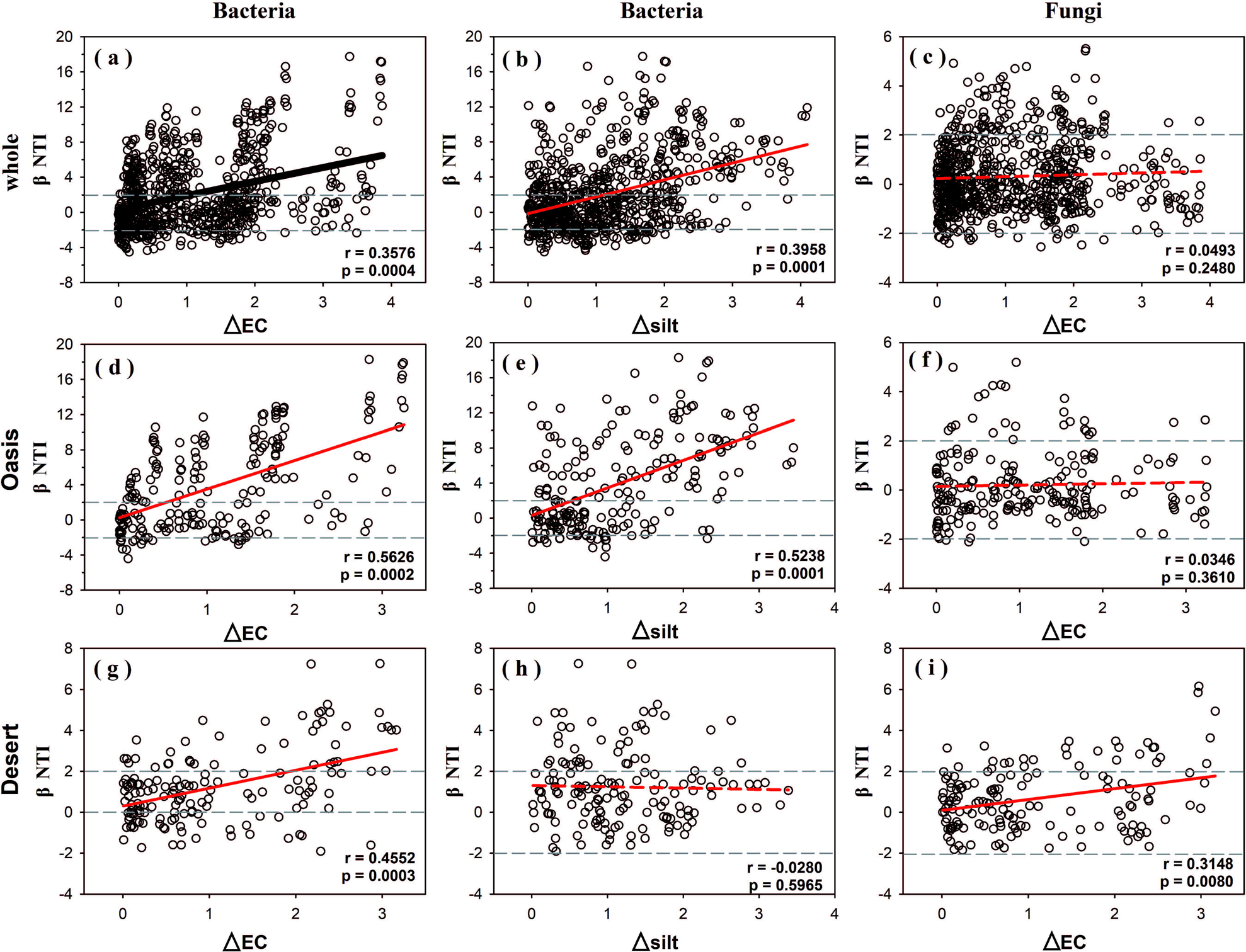
The relationships between βNTI values and differences in EC (a, d, g) and soil silt content (b, e, h) for bacteria and in EC for fungi (c, f, i) in different habitats. Horizontal dashed gray lines indicate upper and lower significant thresholds at βNTI=+2 and βNTI=-2, respectively.

We further examined the role of EC on microbial community assembly by testing whether assembly processes varied with respect to EC levels. Bacterial phylogenetic turnover was dominated by the deterministic processes, especially variable selection (44.4%) in low EC, while stochastic processes, especially dispersal limitation (60.0%), were more prevalent in High EC (Fig. 4a,b). In contrast, stochastic processes dominated fungal community assembly both in Low EC and High EC (96.6% and 94.6%, respectively), especially dispersal limitation (83.6% and 80.0%, respectively; Fig. 4c,d).

**Figure 4.**
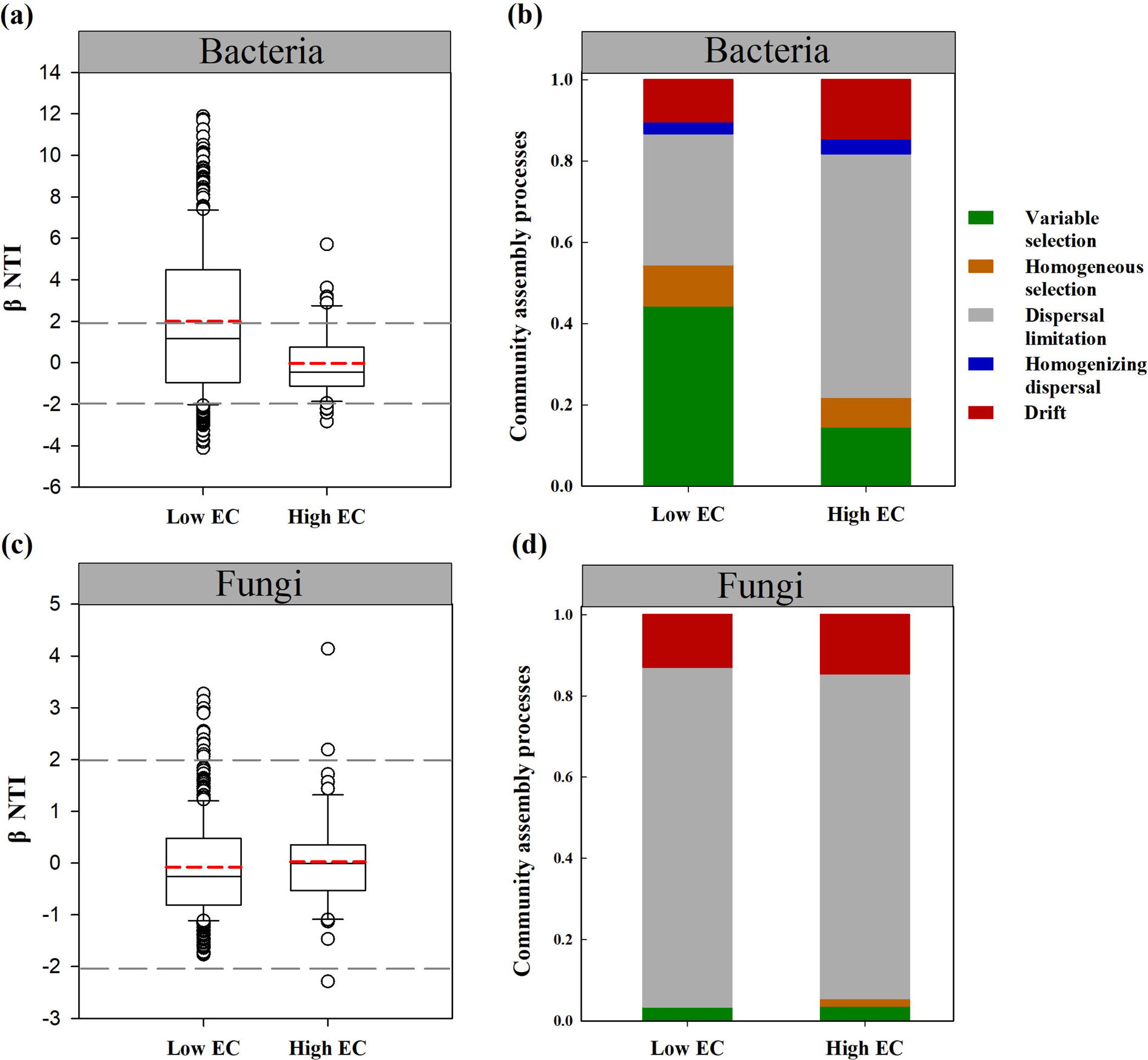
Contributions of deterministic and stochastic processes on bacterial (a, b) and fungal (c, d) community assembly within EC groups. The threshold value dividing low EC and high EC was 8ms/cm.

### Biotic association also play a nonnegligible role on microbial community assembly

In addition to environmental filtering, associations between taxa are also important selection force for microbial community assembly (Cosetta and Wolfe, 2019). Based on cross-kingdom co-occurrence networks, which consisted of 1456 vertexes and 37032 edges (Fig. S4). We found that soil silt and sand content and EC showed the significantly correlations with topological features (Fig. S4). Mantel test indicated that bacterial phylogenetic turnover (βNTI) was significantly correlated with all topological features in oasis habitat while only with Bac_fun and AD index in desert habitat (Table 3). In contrast, fungal βNTI had no significantly relationship with topological features, except for AD in desert habitat (Table 3). Furthermore, results of piecewise SEM showed that both biotic associations and EC had significant influence on bacterial βNTI, and EC would indirectly affect bacterial βNTI through biotic associations (Fig. 5a,b,c). However, most variables had no effects on fungal βNTI, except that EC had significantly influence on fungal βNTI in desert habitat(Fig. 5d,e,f).

**Figure 5.**
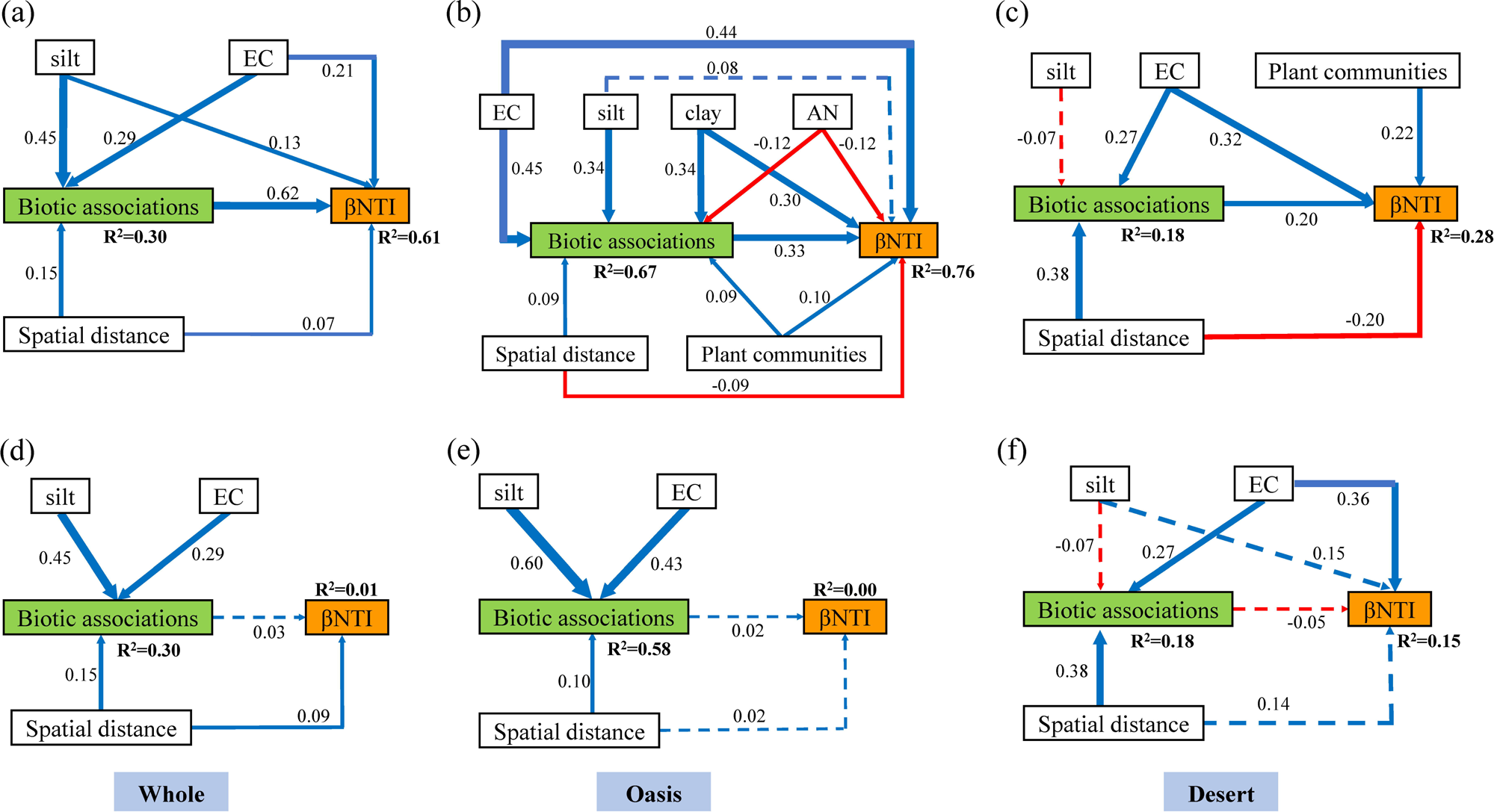
Piecewise structural equation models (piecewise SEM) show the direct and/or indirect effects of soil variables, spatial distance, biotic associations and plant communities on phylogenetic turnover (βNTI) for bacterial (a, b, c) and fungal (d, e, f) communities in different habitats. R2 represent the proportion of variance explained by variables. Solid red and blue arrows indicate negative and positive relationships, respectively, while dashed arrows indicate no significant relationships. EC, soil electrical conductivity; AN, available nitrogen.

**Table 3.**
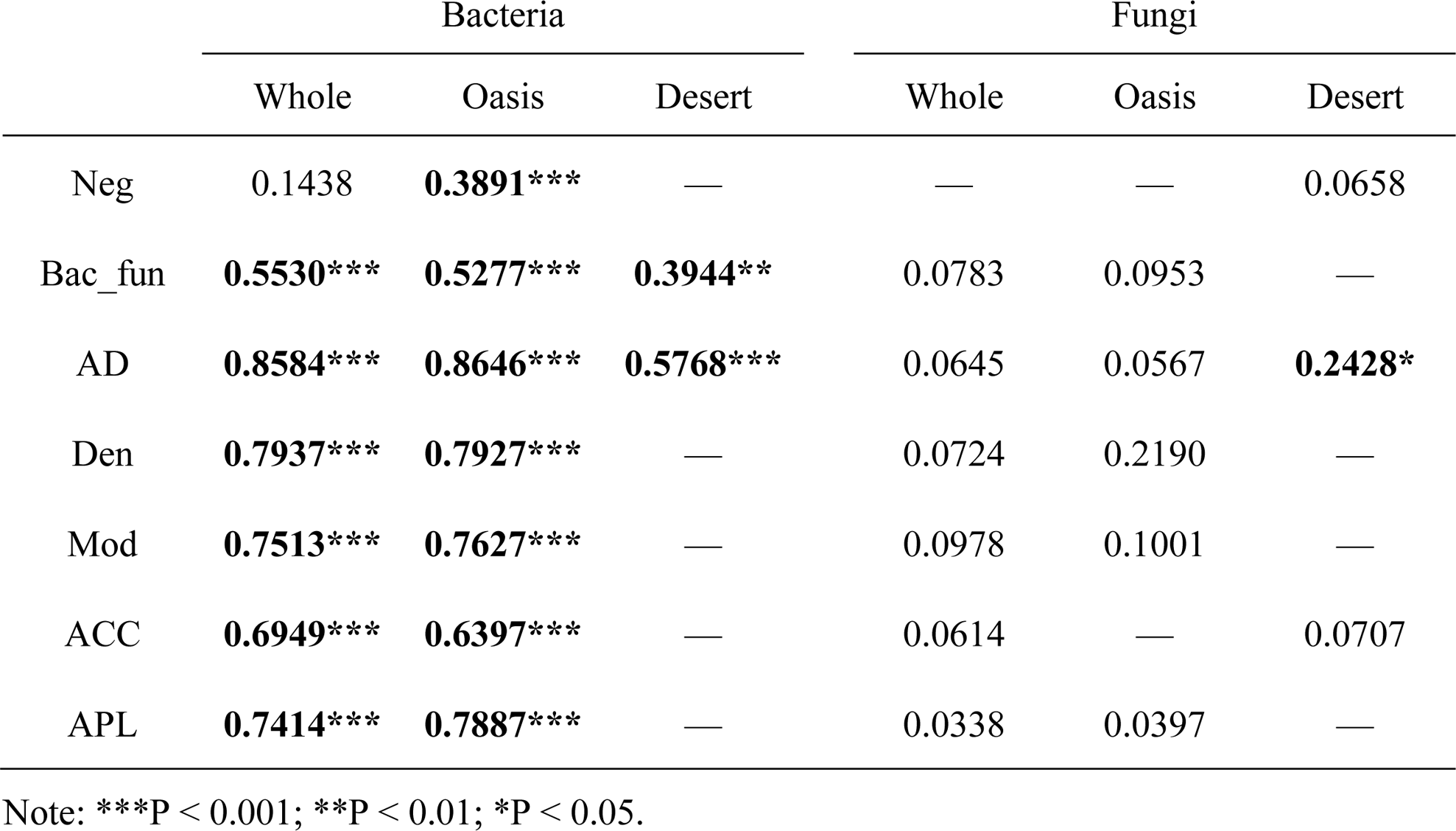
The relationships between phylogenetic turnover (βNTI) of bacterial and fungal communities and indexes of biotic associations based on Mantel test.

|  | Bacteria |  |  | Fungi |  |  |
| --- | --- | --- | --- | --- | --- | --- |
|  | Whole | Oasis | Desert | Whole | Oasis | Desert |
| Neg | 0.1438 | **0.3891\*\*\*** | — | — | — | 0.0658 |
| Bac_fun | **0.5530\*\*\*** | **0.5277\*\*\*** | **0.3944\*\*** | 0.0783 | 0.0953 | — |
| AD | **0.8584\*\*\*** | **0.8646\*\*\*** | **0.5768\*\*\*** | 0.0645 | 0.0567 | **0.2428\*** |
| Den | **0.7937\*\*\*** | **0.7927\*\*\*** | — | 0.0724 | 0.2190 | — |
| Mod | **0.7513\*\*\*** | **0.7627\*\*\*** | — | 0.0978 | 0.1001 | — |
| ACC | **0.6949\*\*\*** | **0.6397\*\*\*** | — | 0.0614 | — | 0.0707 |
| APL | **0.7414\*\*\*** | **0.7887\*\*\*** | — | 0.0338 | 0.0397 | — |

Note: \*\*\*P < 0.001; \*\*P < 0.01; \*P < 0.05.

## Discussion

### Relative importance of the community assembly processes of soil bacteria and fungi in oasis and desert habitats

Our results showed that deterministic processes, especially variable selection, were the main drivers for bacterial community assembly in oasis habitat, while stochastic processes, especially dispersal limitation, dominated the bacterial community assembly in desert habitat (Fig. 2). These results indicated that heterogenous environment exerted stronger selection pressure on bacterial communities in oasis than desert habitat, which was in contrast with previous studies having confirmed that extreme environment (such as extreme acidic or alkaline pH) would impose strong filtering effects for specific microorganisms so as to result in dominance of deterministic processes (Valverde *et al*., 2014; Tripathi *et al*., 2018). The possible reason is that variations of soil environment was greater in oasis habitat than desert habitat (Wang *et al*., 2019). The small environmental variations would lead to decrease of variable selection yet increasing contribution of stochastic processes on community assembly (Wang *et al*., 2013).

In contrast to bacteria, fungal community assembly was dominated by stochastic processes, especially dispersal limitation, both in oasis and desert habitats (Fig. 2). This finding is similar to previous study in the Thousand Island Lake (Wang *et al*., 2020) and Qinling Mountains (Huo *et al*., 2023), showing that stochastic processes dominated fungal community assembly while deterministic processes dominated bacterial community assembly. These results indicated that fungal community assembly processes maybe dominated by stochastic processes in most environments or ecosystems. In addition, in oasis habitat, fungal community assembly was largely dominated by dispersal limitation (83.8%) while bacterial community assembly was mainly affected by variable selection (Fig. 2). This finding was consistent with previous work showing that bacterial but not fungal community turnover were significantly influenced by environmental changes (Rousk *et al*., 2010; Powell *et al*., 2015). One possible explanation for the dominance of dispersal limitation for fungal communities is owing to the relatively larger “body size” of fungi (3-21μm), which would limit their long-distance dispersal compared to the smaller-sized bacteria (0.5-10μm) (Bässler *et al*., 2015). In addition, extending mycelia of fungi can promote their absorption of water or nutrients, and this behavior would make them more resistant to variations of environment (Guhr *et al*., 2015), thereby reducing the influence of variable selection on fungal community assembly.

### Factors influencing community assembly processes of soil bacteria and fungi

Some recent studies have identified the influence of soil pH and soil salinity on soil bacterial community assembly (Tripathi *et al*., 2018; Zhang *et al*., 2019), while our study emphasized the selection force of soil EC and soil particle composition on soil bacterial community assembly (Table 2). As the △EC increased, bacterial community assembly altered from stochastic to deterministic processes, whereas △EC was not enough to change fungal community assembly processes (Fig. 3). The possible explanation for this different effect could be attributed to higher tolerance of fungi than bacteria to salinity (Rath *et al*., 2019a). chitinous cell walls of fungi can provide protection against low matric potentials and their mycelia networks enable translocation and redistribution of soil water (Guhr *et al*., 2015).

Soil characteristics are known to impose selection pressures on microbial community, but which vary at different levels of each soil properties (Dini-Andreote *et al*., 2015; Tripathi *et al*., 2018). For instance, a previous study has reported that extreme acidic or alkaline pH conditions lead to dominance of deterministic processes, whereas neutral pH conditions lead to more stochastic assembly processes (Tripathi *et al*., 2018). In contrast, our study confirmed that extremely high EC lead to dominance of stochastic processes in bacterial community assembly, but lower EC level lead to stronger deterministic processes (Fig. 4). It was consistent with previous studies with a low EC ranges (0ms/cm ∼ 2.5ms/cm) that showed deterministic processes have overtaken stochastic processes in microbial community assembly (Zhang *et al*., 2019). Such results might be contributed to less sensitive of microbial communities to EC increases in already highly EC soils than low EC soils (Rath *et al*., 2019b). Additionally, extremely high EC may inhibit environmental filtering exerted by other factors, resulting in a decrease in the contribution of deterministic processes.

Fungal community assembly was dominated by dispersal limitation, and environmental variables showed little/no explanation on fungal βNTI (Table 1 & Table 2). However, our previous study confirmed that plant attributes had significantly influence on fungal community turnover (Wang *et al*., 2021a). The possible explanation was that plant community may naturally form the “plant barriers” resulting in lower dispersal rates of fungi, which would increase the contribution of dispersal limitation. According to a previous study, undegraded meadow had more contribution of dispersal limitation on fungal community assembly than degrade meadow, suggesting that plants could decrease fungal dispersal rate (Wang *et al*., 2021b). In addition, we found that effect of dispersal limitation on fungal community assembly was greater in oasis habitat (83.8%) than desert habitat (58.8%), which further demonstrate the function of “plant barriers” on account of higher vegetation coverage in oasis habitat than desert habitat.

Our results showed that networks’ topology features/biotic associations were significantly correlated with bacterial βNTI, but not fungal βNTI (Fig. 5), which further confirmed the dominance of stochastic processes on fungal community assembly in arid inland river basin. In addition, networks’ topology was differently correlated with environmental factors, which complicated the relationship between environmental factors, biotic interactions and microbial assembly processes. For instance, soil EC was negatively correlated with AD while positively related with APL (Fig. S4), suggesting that extreme EC may weaken the connections among microorganisms and further indirectly affect microbial community assembly (Fig. 5).

## Conclusions

In summary, bacterial community assembly were dominated by deterministic processes in oasis habitat whereas that were dominated by stochastic processes in desert habitat. In contrast, stochastic processes were major mechanism driving fungal community assembly in both oasis and desert habitats. Soil EC play an important role in regulating the relative contributions of deterministic and stochastic processes on bacterial community assembly in arid inland river basin. Besides, the contribution of biotic interactions could also not be ignored. However, both environmental factors and biotic interactions had little/no effects on fungal community assembly. This study suggests that assembly processes of soil microbial community differed depending on the habitat and taxa.

## Acknowledgements

We would like to gratefully thank the Funding of this work that was granted by the National Natural Science Foundation of China (32271703, 31971538), Hainan University Research Start-up Fund (KYQD(ZR)-21110) and Hainan Provincial Natural Science Foundation of China (322QN247). We are also grateful to Guorui Xu in Xishuangbanna Tropical Botanical Garden, Chinese Academy of Sciences for his suggestions on the manuscript.

## Author contributions

Wenjuan Wang and Jingwen Li designed the study and collected the data. Wenjuan Wang, Yiming Ding and Jingwen Li analyzed the data. Wenjuan Wang wrote the first draft of the manuscript, Jingwen Li and Mingxun Ren commented on the manuscript. All authors read and approved the final manuscript.

