## supplementary for "Assembly processes of bacterial and fungal community differ between desert and oasis habitats in an arid inland river basin, northwest China"

Corresponding author*:

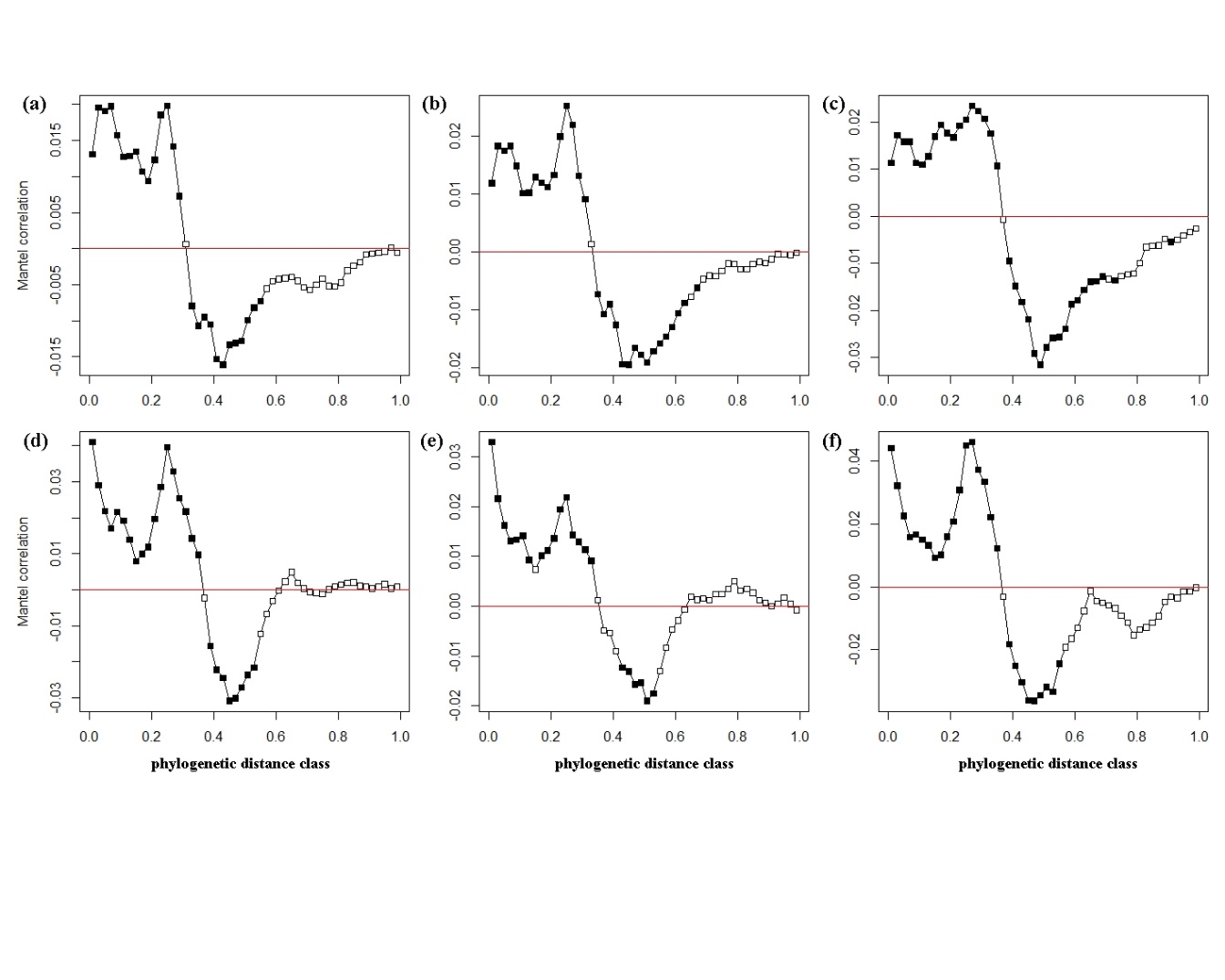

(Whole)

(Whole)

(Oasis)

(Desert)

(Oasis)

(Desert)

**Figure S1** Mantel correlograms between the pairwise matrix of OTU’s niche distances and phylogenetic distance for bacteria (a-c) and fungi (d-f). Closed squares indicate significant correlations (P<0.05) after Bonferroni correction for multiple tests.

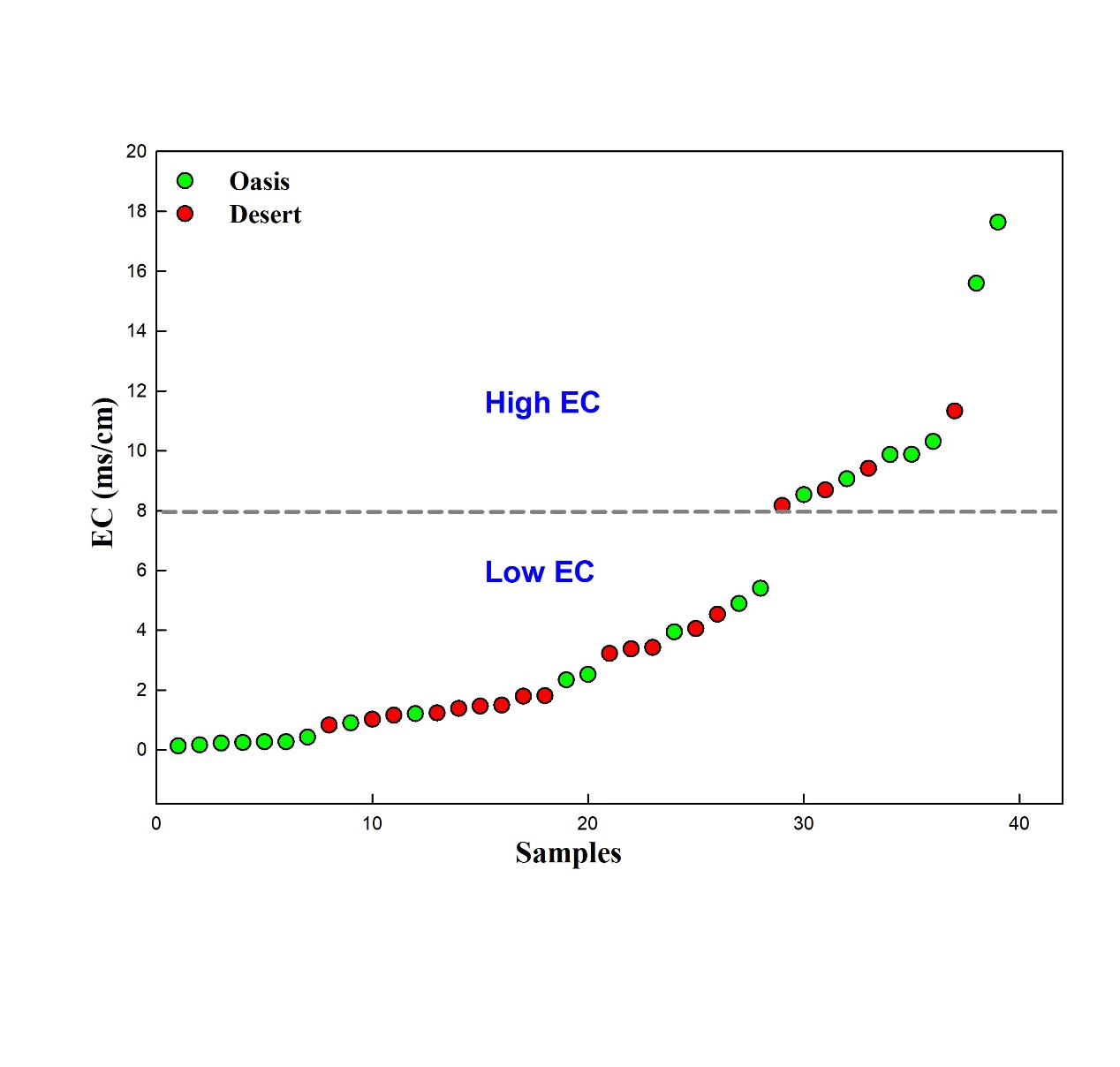
**Figure S2** Scatter plot of soil electrical conductivity (EC) of all samples. The dashed grey line represents threshold value (8ms/cm) dividing low EC and high EC groups.

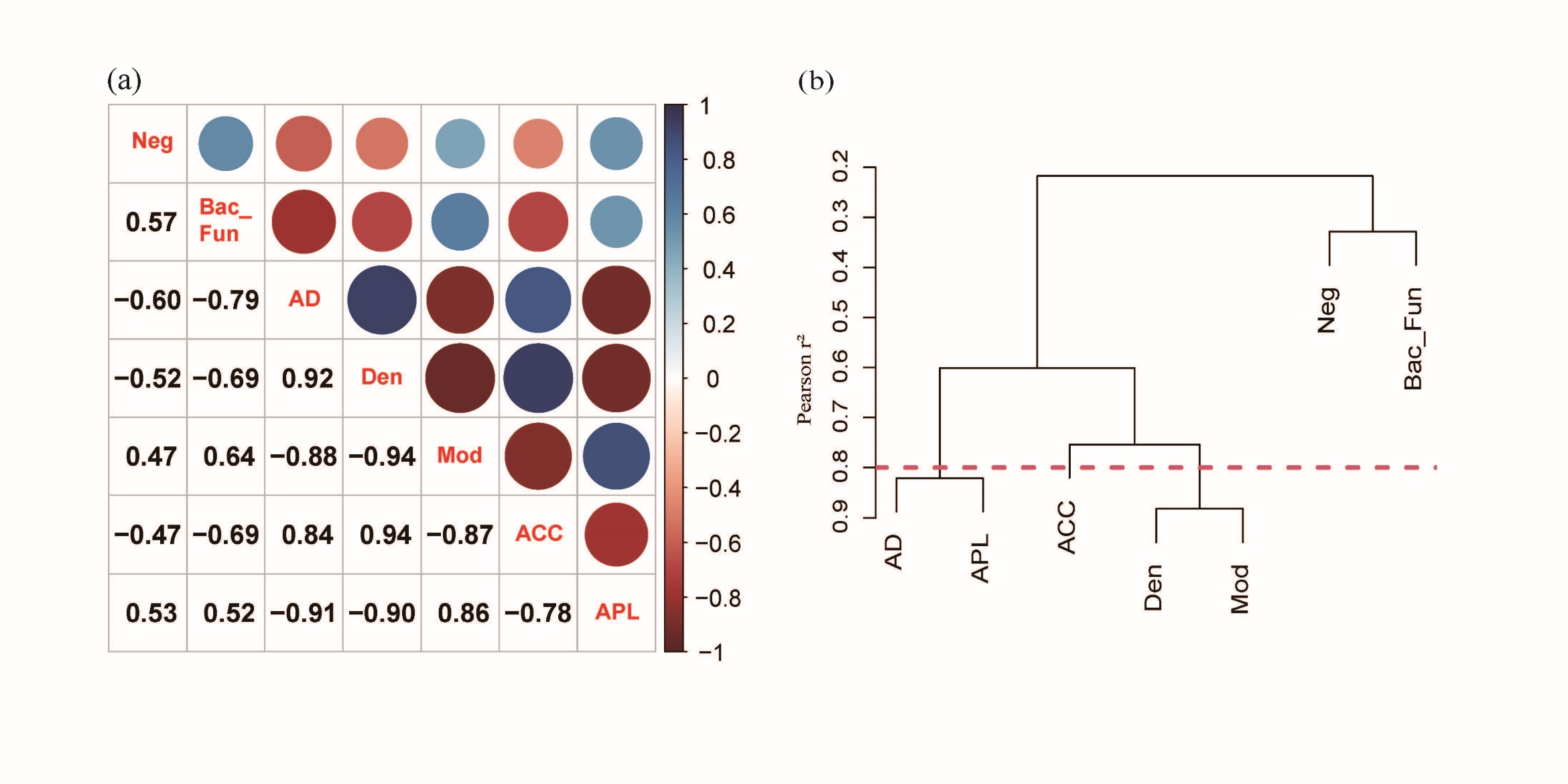

**Figure S3** Correlations of indexes of biotic associations within ecological network. (a) Correlations heatmap of indexes of biotic associations within ecological network. (b) cluster analysis of indexes of biotic associations within ecological network.

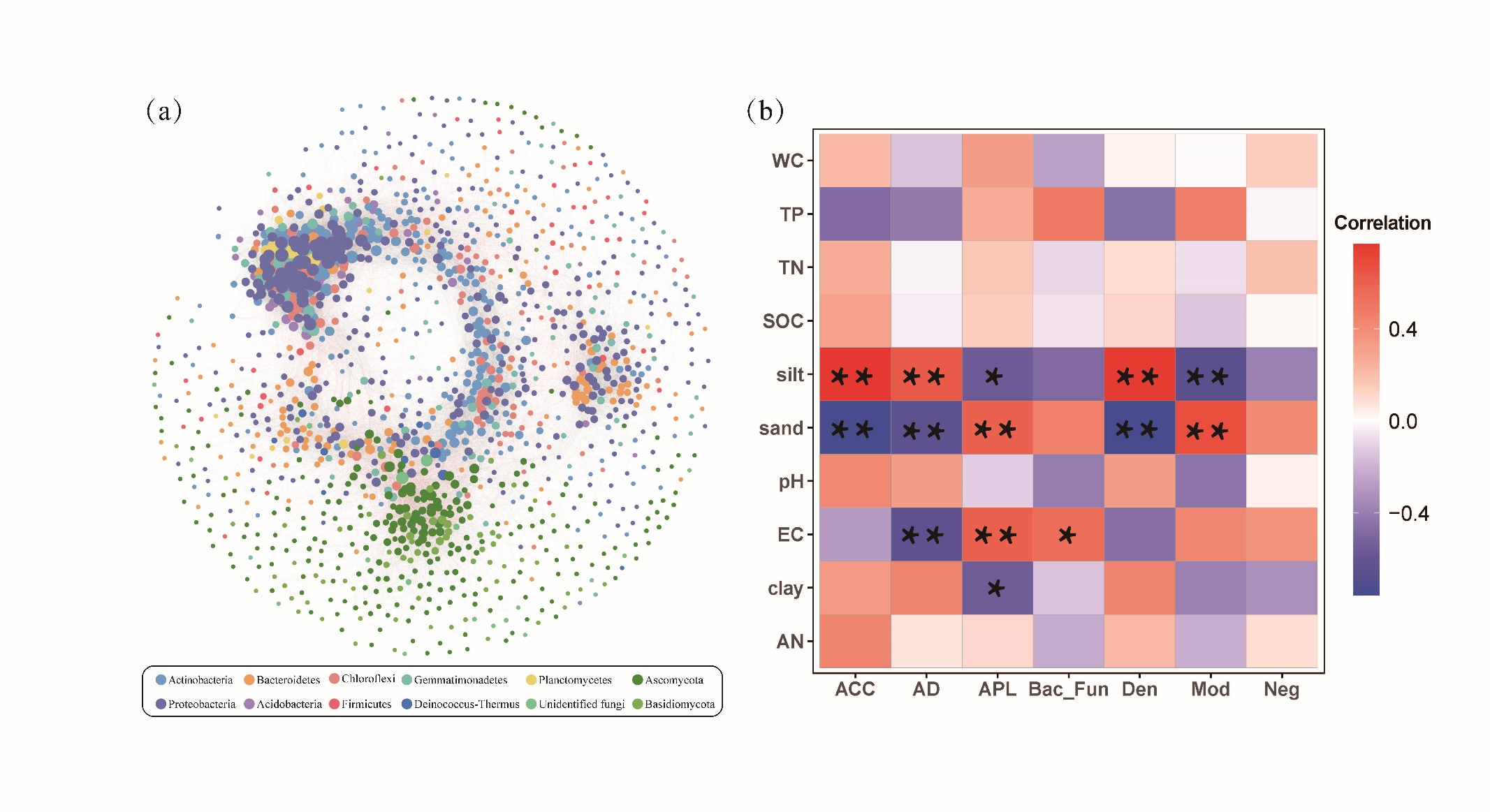

**Figure S4** Co-occurrence network of microbial community (including bacteria and fungi) and its relationships with soil variables. (a) Cross-kingdom co-occurrence network of microbial taxa. The vertexes are colored based on the taxa phylum. The size of vertexes is proportional to the degree of the taxa. The connection indicates a strong and significant (P<0.001) correlation, while light grey and red edges represent positive (Spearman’s ρ > 0.7) and negative (Spearman’s ρ < -0.7) correlations, respectively. (b) Correlations between indexes of biotic associations within ecological network and soil variables. ACC, average clustering coefficient; AD, average degree; APL, average path length; Bac_Fun, the proportion of interacted associations between bacterial and fungal taxa; Den, density of the network; Mod, modularity; Neg, the proportion of negative associations.
